# Human sensory-like neuron cultivation – an optimized protocol

**DOI:** 10.1101/2024.06.10.598264

**Authors:** Nicole Michelle Schottmann, Julia Grüner, Frederik Bär, Franziska Karl-Schöller, Sabrina Oerter, Nurcan Üçeyler

## Abstract

**Introduction:** Reprogramming of human induced pluripotent stem cells (iPSC) and their differentiation into specific cell types, such as induced sensory-like neurons (iSN), are critical for disease modeling and drug testing. However, the variability of cell populations challenges reliability and reproducibility. While various protocols for iSN differentiation exist, development of non-iSN cells in these cultures remains an issue. Therefore, standardization of protocols is essential. This study aimed to improve iSN culture conditions by reducing the number of non-iSN cells while preserving survival and quality of iSN.

**Methods:** iSN were differentiated from a healthy control iPSC line using an established protocol. Interventions for protocol optimization included floxuridine (FdU) or 1-β-D-arabinofuranosyl-cytosin-hydrochlorid (AraC) treatment, Magnetic-Activated Cell Sorting (MACS), early cell passaging, and replating. Cell viability and iSN-to-total-cell-count ratio were assessed using a luminescent assay and immunocytochemistry, respectively.

**Results:** Passaging of cells during differentiation did not increase the iSN-to-total-cell-count ratio, and MACS of immature iSN led to neuronal blebbing and reduced the iSN-to-total-cell-count ratio. Treatment with high concentrations and prolonged incubation of FdU or AraC resulted in excessive cell death. However, treatment with 10 μM FdU for 24 h post-differentiation showed the most selective targeting of non-iSN cells, leading to an increase in the iSN-to-total-cell count ratio without compromising the viability or functionality of the iSN population. Replating of iSN shortly after seeding also helped to reduce non-iSN cells.

**Conclusion:** In direct comparison to other methods, treatment with 10 μM FdU for 24 h after differentiation represents a promising approach to improve iSN culture purity, offering a potential benefit for downstream applications in disease modeling and drug discovery. However, further investigations involving multiple iPSC lines and optimization of protocol parameters are warranted to fully exploit the potential of this method and enhance its reproducibility and applicability. Overall, this study provides valuable insights into optimizing culture conditions for iSN differentiation and highlights the importance of standardized protocols in iPSC-based research.

## Introduction

Generation of human induced pluripotent stem cells (iPSC) (Takahashi and Yamanaka, 2006) and their differentiation into specific target cells such as sensory-like neurons (iSN) (Chambers et al., 2009) has developed into a potent method of disease modeling and drug testing. Standardization of methodological procedures is crucial to reduce technical variability to a minimum and to ensure reliability and reproducibility (Lampert et al., 2020; Volpato and Webber, 2020). To date, two protocols are available for differentiating iSN, namely based on combined small-molecule inhibition (Chambers et al., 2012) and on overexpression of transcription factors (Blanchard et al., 2015). Application of the small-molecule protocol also results in the generation of non-iSN cells with high variability in morphology and counts between differentiations (Schwartzentruber et al., 2018). This cellular heterogeneity challenges correct data allocation and interpretation.

To increase culture purity, the number of non-neuronal cells can be reduced by the application of cytostatic compounds used in chemotherapy (Hilgenberg and Smith, 2007; Thirumangalakudi et al., 2009; Irobi et al., 2010; Liu et al., 2013; Schwieger et al., 2016; Clark et al., 2021). This inhibits proliferation of non-iSN by mainly targeting mitotic cells. However, no standardized protocol is available as for the best compounds, time points, and concentrations (Hilgenberg and Smith, 2007; Thirumangalakudi et al., 2009; Irobi et al., 2010; Liu et al., 2013; Schwieger et al., 2016; Clark et al., 2021). Further approaches achieved higher ratios of iSN-to-non-iSN cells by additional splits during the first days of iSN differentiation (Clark et al., 2017) and neuronal cell sorting (Hirano et al., 2021).

We aimed to improve culture conditions of human iSN by reducing the number of non-iSN cells in culture, while preserving iSN quality. We assessed cell viability and iSN-to-total-cell count ratios after testing defined *in vitro* interventions. Among these, we found that the application of the chemotherapeutic floxuridine (FdU) 10 μM for 24 h after differentiation yielded the best results. We demonstrate that this treatment improves culture purity by being the most selective in targeting non-iSN cells without compromising the long-term viability or functionality of the iSN population. FdU led to an increase in the iSN-to-total-cell count ratio indicating a reduction of non-iSN cells. We further show that these iSN express sensory neuron-specific marker proteins and are electrically active.

## Materials and methods

### iSN differentiation

iSN were differentiated from a healthy control iPSC line as described recently (Klein et al., 2024). In brief, 1.125*10^6^ iPSC were seeded into Growth Factor Reduced Matrigel-coated wells (bMg, Corning, Corning, NY, USA) in a six-well plate in 2 ml StemMACS iPS-Brew (Miltenyi Biotec, Bergisch Gladbach, Germany). Medium was supplemented with 100 U/ml penicilin/streptomycin (Gibco, Waltham, MA, USA) and 10 μM Y27632 (Miltenyi Biotec, Bergisch Gladbach, Germany) on day −2. On day −1, StemMACS medium was changed and differentiation was started on day 0 after passaging. Cells were then cultivated in KnockOut medium (KSR; KnockOut DMEM/F12, 2 mM GlutaMAX, 15% KnockOut Serum Replacement, 100 μM 2-mercaptoethanol, 0.1 mM minimum essential medium non-essential amino acids + 100 U/ml penicilin/streptomycin [all: Thermo Fisher Scientific, Waltham, MA, USA]). The KnockOut medium was supplemented with a two-inhibitor cocktail (2i) containing 100 nM LDN-193189 (Stemcell Technologies, Vancouver, Canada) and 10 μM SB-431542 (Miltenyi Biotec, Bergisch Gladbach, Germany). Three inhibitor (3i) cocktail (10 μM SU-5402, 10 μM DAPT [both: Sigma Aldrich, St. Louis, MO, USA]), and 3 μM CHIR-99021 (Axon Medchem, Groningen, Netherlands) was added from day +2. Starting on day +4, KSR medium was replaced in 25%-steps with N2 medium (N2; DMEM/F12 GlutaMAX, 1X B-27 Plus Supplement, 1X N-2 Supplement, 100 U/ml penicillin/streptomycin [all: Thermo Fisher Scientific, Waltham, MA, USA]) every other day. Cells were passaged on bMg-coated 12-mm coverslips with a 1:2 ratio on day 10 using TrypLE Express (Thermo Fisher Scientific, Waltham, MA, USA) for 8-20 min. If not stated otherwise, neuronal maturation medium consisted of N2 medium supplemented with 20 ng/ml BDNF, 20 ng/ml GDNF, 20 ng/ml NGFb (all: Peprotech, Rocky Hill, NJ, USA), and 200 ng/ml ascorbic acid (Sigma-Aldrich, St. Louis, MO, USA). iSN matured for 6 weeks.

### FdU treatment

iSN were seeded on bMg-coated coverslips on day +10 of differentiation and were treated with FdU (Santa Cruz Biotechnology, Dallas, TX, USA) 1 μM, 5 μM, 10 μM, and 20 μM for 24 h, 48 h, and 72 h. For later time points, half medium change was performed to ensure stable FdU concentrations. Results were compared to 10 μM FdU for 24 h as a control (ctrl).

### 1-β-D-arabinofuranosyl-cytosin-hydrochlorid treatment

iSN were seeded on bMg-coated coverslips on day +10 of differentiation. Cells were incubated with 0.5 μM, 1 μM, 5 μM, and 10 μM of 1-β-D-arabinofuranosyl-cytosin-hydrochlorid (AraC, Sigma Aldrich, St. Louis, MO, USA) over 24 h, 48h, 72 h, 120 h, and 168 h. Stable AraC concentrations were ensured by performing half media changes for longer incubation times. As a control condition, cells were treated with 10 μM FdU for 24 h.

### Magnetic Activated Cell Sorting

Following a published protocol (Hirano et al., 2021), iSN were sorted using Magnetic Activated Cell Sorting (MACS) on day +10 of differentiation. Cells were detached, resuspended in 1 ml phosphate buffered saline with Mg2+ and Ca2+ (PBS ++, PBS, Merck, Darmstadt, Germany), and counted as described in iSN differentiation. Cells were sorted using Neural Crest Stem Cell MicroBeads, human (Miltenyi Biotec, Bergisch Gladbach, Germany). Afterwards, cells were seeded on bMg-coated coverslips in 1 ml maturation medium, additionally containing 10 μM DAPT. Two further approaches were tested: A cell strainer was used to reduce cell clusters as well as a second MACS step was included for higher purification. Cells incubated with 10 μM FdU for 24 h were used as control.

### Split on day +2 of differentiation

Cells were splitted on day +2 of differentiation following Clark et al. [14]. Cells were detached after washing with PBS using 0.5 mM EDTA (Invitrogen, Carlsbad, CA, USA) for 5 min at 37°C. Cells were splitted 1:1 onto bMg-coated wells in 6-well plates containing KSR medium supplemented with 10 μM Y27632 and LDN, SB, CHIR, DAPT, and SU as described in iSN differentiation.

### Replating of iSN

iSN were replated on freshly bMG-coated coverslips after 10, 15, 20, 25, 30, and 45 min following day +10 split. Media was directly transferred on a new coverslip without further resuspension. Old coverslips were substituted with fresh media each.

### Immunocytochemistry

Cells were fixed using 4% paraformaldehyde (PFA; Electron Microscopy Sciences, Hatfield, USA) for 20 min at room temperature followed by three washing steps with PBS++. Blocking solution was applied for 30 min at room temperature and consisted of 10% fetal bovine serum (Honduras Origin, Sigma Aldrich, St. Louis, MO, USA) and 0.1% Saponin (Sigma Aldrich, St. Louis, MO, USA) diluted in PBS++. Anti-voltage-gated sodium channel 1.8 (Nav1.8) antibody (1:100, Abcam, Cambridge, UK), peripherin (PRPH) antibody (1:250, AF488/AF647, Santa Cruz Biotechnology, Dallas, TX, USA), and anti-beta-III-tubulin (TUJ1) antibody (1:500, Abcam, Cambridge, UK) diluted in blocking solution were applied overnight at 4°C. Unbound antibody was washed with PBS++ including 4’,6-diamidino-2-phenylindole for initialization (DAPI; 1:10 000, Sigma Aldrich, St. Louis, MS, USA). Where needed, secondary antibodies (donkey anti-chicken AF488 and donkey anti-rabbit Cy3; Jackson ImmunoResearch Europe, Cambridge House, St. Thomas’ Place, Ely, UK) were applied for 30 min after washing of primary antibodies three times with PBS++ and before adding DAPI. Cells were mounted in Aqua-Poly/Mount (Polysciences, Warrington, PA, USA). Photomicrographs for further analysis were taken with an inverted fluorescence microscope (Leica DMi 8, Leica Microsystems, Wetzlar, Germany). Analysis was performed using a CellProfiler 4.1.3. Ink pipeline. Other images were captured using a Zeiss Axio Imager M2 microscope.

### Cell viability assay

ISN were seeded on bMg-coated 96-well plates (Greiner Bio-One GmbH, Frickenhausen, Germany) on day +10 of differentiation, and incubated with AraC or FdU as described above. After 24, 48, and 72 h, a CellTiter-Glo® Luminescent Cell Viability Assay (Promega, Madison, WI, USA) was performed. Cells received 1:1 maturation medium and CellTiter-Glo® Reagent (Promega, Madison, WI, USA). Plates were shaken at 480 rpm for 2 min followed by 10 min incubation at room temperature in the dark. Luminescence was measured using a Tecan Spark multiplate reader (Tecan Trading AG, Männedorf, Switzerland).

### Multi-electrode array

Electrical activity of iSN was assessed by a Multiwell multi-electrode array (MEA) system (Multi Channel Systems MCS GmbH, Reutlingen, Germany). ISN were seeded into bMg-coated 24-well plates with PEDOT electrodes on glass (Multi Channel Systems, Reutlingen, Germany) on day +10 of differentiation in neuronal maturation medium supplemented with 10 μM FdU. MEA measurements were performed after 4, 5, and 6 weeks of maturation, respectively.

## Results

### Passaging of cells during differentiation does not increase iSN-to-total-cell-count ratio

Passaging of cells on day +2 of differentiation did not result in morphological changes as shown in representative bright field (Fig. 1A) and ICC photomicrographs (Fig. 1B). Analysis of ICC images via a cell profiler pipeline did not reveal any differences in the iSN-to-total-cell-count ratio between cells splitted on day +2 of differentiation and ctrl (Fig. 1C). While the ratio did not change, the number of generated cells in general increased (p < 0.001; Fig. 1D).

**Figure 1:**
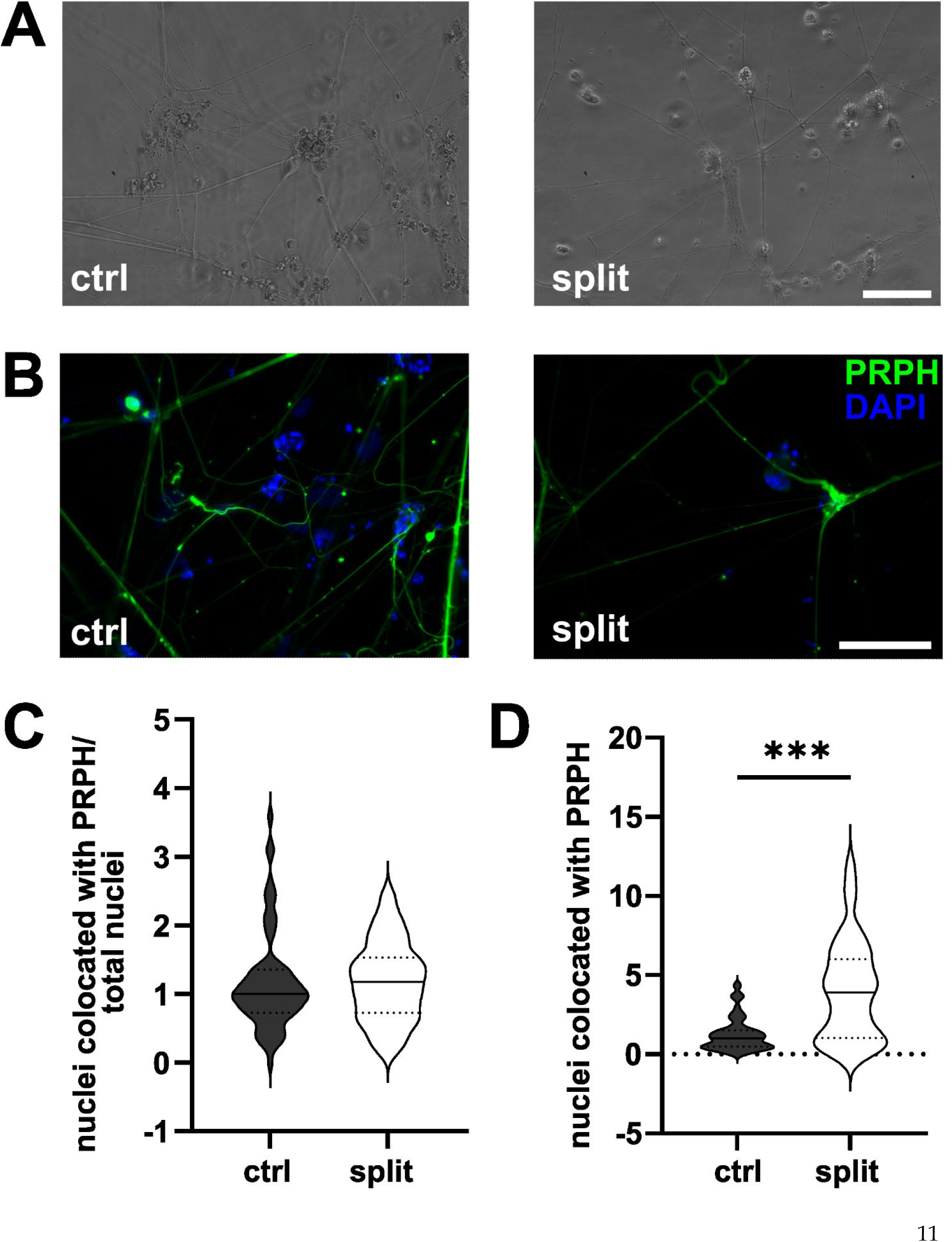
iSN splitted on day +2 of differentiation. Representative bright field (A) and ICC photomicrographs (B) of splitted cells compared to ctrl. While ratio of iSN-to-total-cell-count remained similar between splitted and ctrl cells (C), the total number of cells produced could be improved by split on day +2 of differentiation (D). Data are presented as violin plots, unpaired t-test. *p < 0.05; **p < 0.01; ***p < 0.001. Abbreviations: ctrl = control; ICC = immunocytochemistry; PRPH = peripherin. Scale bar = 100 μm.

### Sorting of cells on day +10 of differentiation does not increase the ratio of iSN-to-total-cell-count

MACS of immature iSN on day +10 of differentiation led to neuronal blebbing (Fig. 2A, B). Comparison of defined MACS strategies showed a reduced iSN-to-total-cell-count ratio in iSN treated with a combination of MACS and cell strainer compared to ctrl (p < 0.01; Fig. 2C). Single and double MACS application resulted in iSN-to-total-cell-count ratios comparable to ctrl (Fig. 2C). In all cases, application of MACS led to reduced number of iSN and flow-through mostly containing other cell types than iSN.

**Figure 2:**
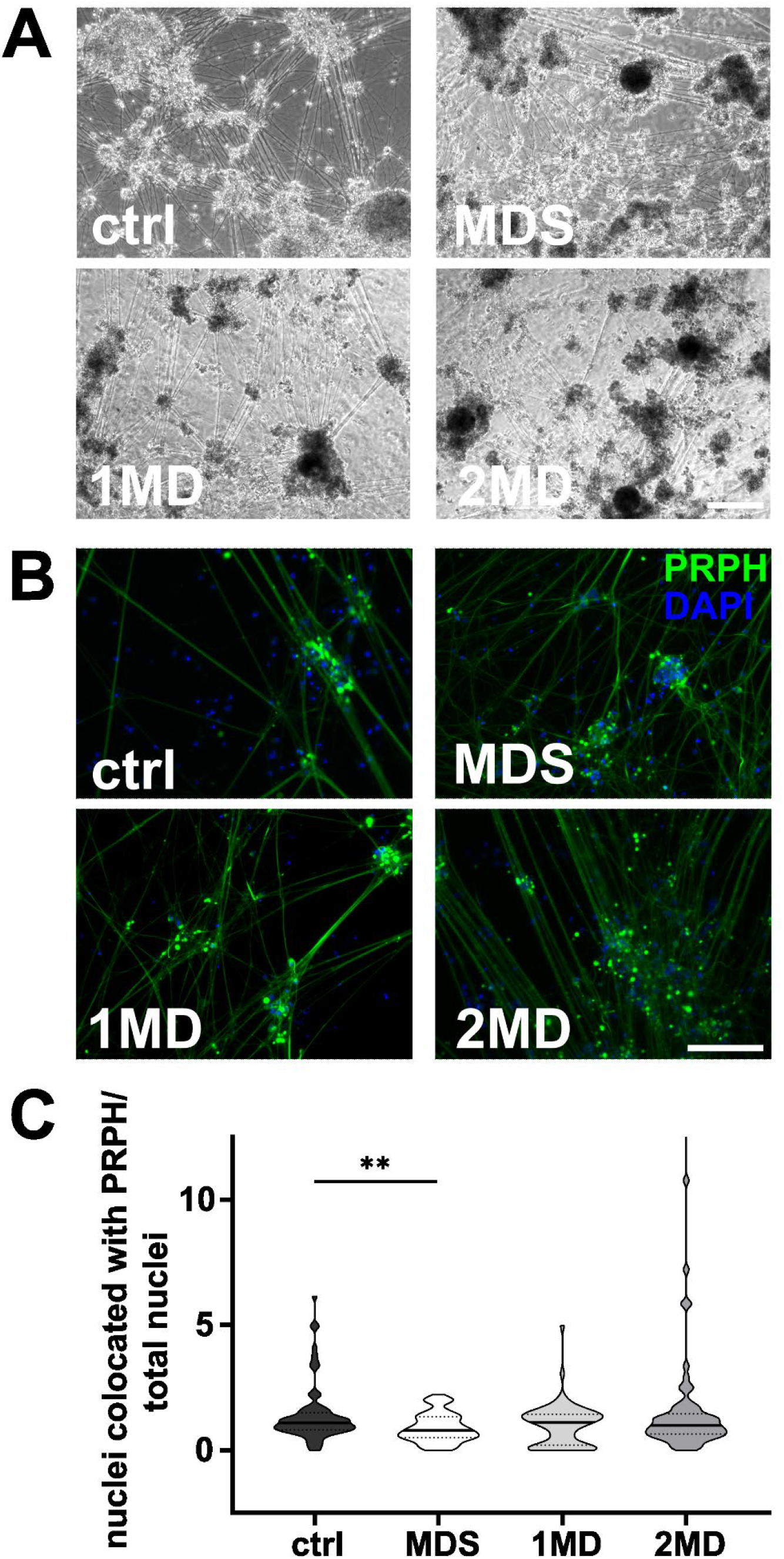
Sorting of iSN on day +10 of differentiation. Representative bright field microscopy (A) and ICC photomicrographs (B) show increased neuronal blebbing of iSN after MACS treatment. Furthermore, non-neuronal cell types changed to a more homogeneous morphology. (C) Combining MACS with the use of a cell strainer resulted in a smaller iSN-to-total-cell-count ratio while single and double MACS applications resulted in a ratio comparable to ctrl. Data are presented as violin plots, Kruskal-Wallis test with Dunn’s multiple comparison. *p < 0.05; **p < 0.01; ***p < 0.001. Abbreviations: ctrl = control; ICC = immunocytochemistry; MD = MACS and DAPT treatment; MDS = MACS and DAPT treatment combined with cell strainer; PRPH = peripherin. Scale bar = 100 μm.

### High concentrations and long FdU or AraC incubation result in excessive cell death

ISN treated with 20 μM FdU did not survive, independent of the duration of incubation. This was also observed when using 10 μM for > 48 h. While incubation with 5 μM FdU > 72 h did not result in complete cell death, only few iSN survived showing morphological signs of stress, e.g., neuronal blebbing (Fig. 3A, B). Therefore, experiments were only repeated for lower concentrations. Application of 1 μM FdU for 24 h was not as effective as the control condition in reducing non-iSN cells (Fig. 3C): also with longer incubation time of 48 h and 72 h, iSN-to-total-cell-count ratio remained smaller (p < 0.01) compared to ctrl. Incubation with 5 μM FdU > 48 h did not increase culture purity (p < 0.001). Only 5 μM applied for 24 h resulted in an iSN-to-total-cell-count ratio similar comparable to ctrl, but trending to a smaller ratio (Fig. 3C), microscopic observation revealed a more pronounced non-iSN layer in most wells.

**Figure 3:**
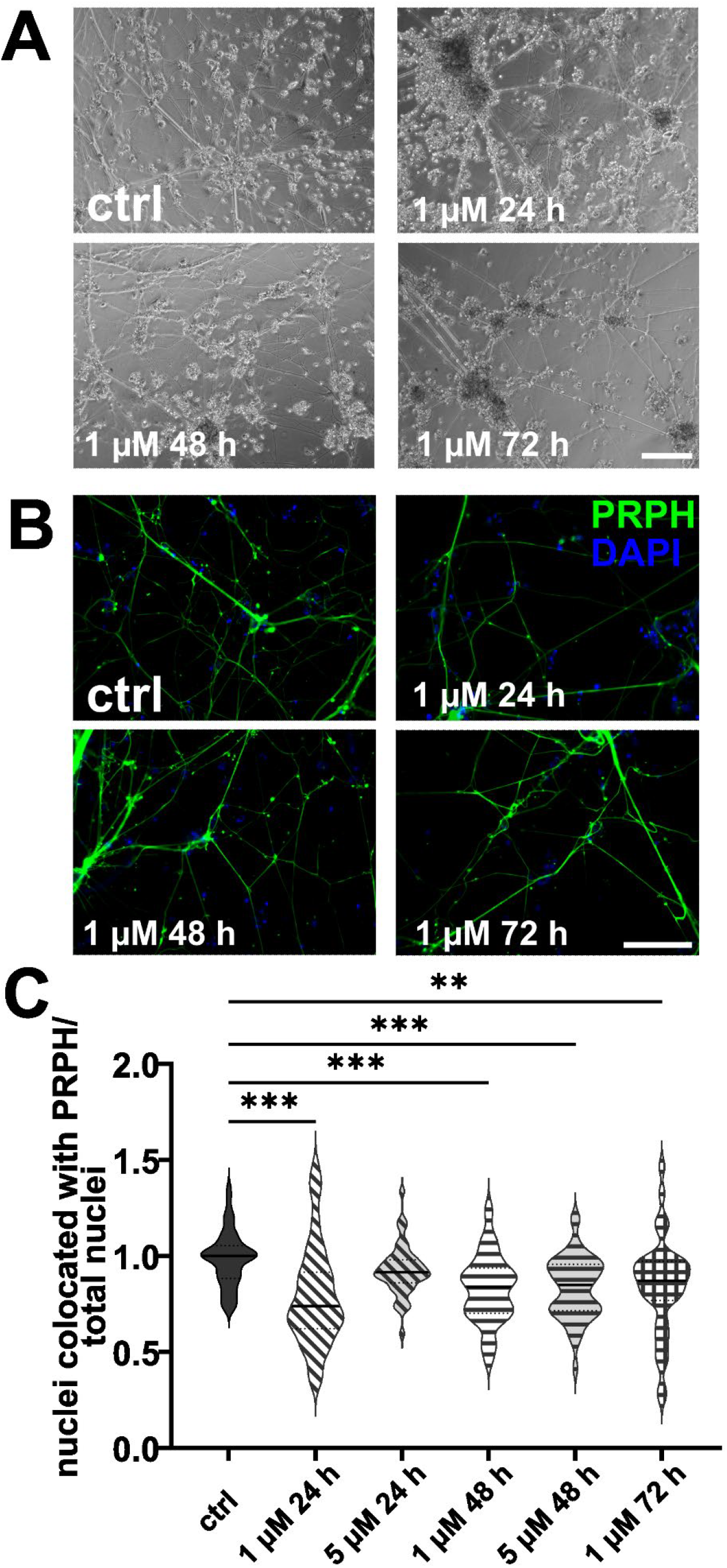
iSN treated with FdU on day +10 of differentiation. Bright field microscopy (A) and ICC photomicrographs (B) did not show an improvement of iSN morphology and reduced number of other cell types with 1 μM FdU over distinct time points (24 h, 48 h, 72 h) compared to ctrl. Most conditions where cells survived showed a smaller ratio of iSN-to-total-cell-count ratio in comparison to ctrl. Application of 5 μM FdU for 24h results in an iSN-to-total-cell-count ratio close to the ctrl (C). Data are presented as violin plots, Kruskal-Wallis test with Dunn’s multiple comparison. *p < 0.05; **p < 0.01; ***p < 0.001. Abbreviations: ctrl = control; ICC = immunocytochemistry; PRPH = peripherin. Scale bar = 100 μm.

In analogy, excessive cell death was observed in iSN treated with 5 μM or 10 μM AraC independent of the incubation time. In contrast, after application of 1 μM AraC for 24 h and 72 h, neuronal cell survival increased. However after longer incubation times, >72 h, iSN completely died. Hence, experimental conditions were adjusted to a lower concentration and shorter incubation time. There was no positive effect on iSN morphology (Fig. 4A, B) and counts of non-iSN cells when treated with 0.5 μM AraC for all time points (Fig. 4C). ISN-to-total-cell-count ratio decreased (p < 0.01) for all treatment conditions compared to ctrl, except for 0.5 μM AraC for 72 h which showed a similar efficacy as ctrl (Fig. 4C).

**Figure 4:**
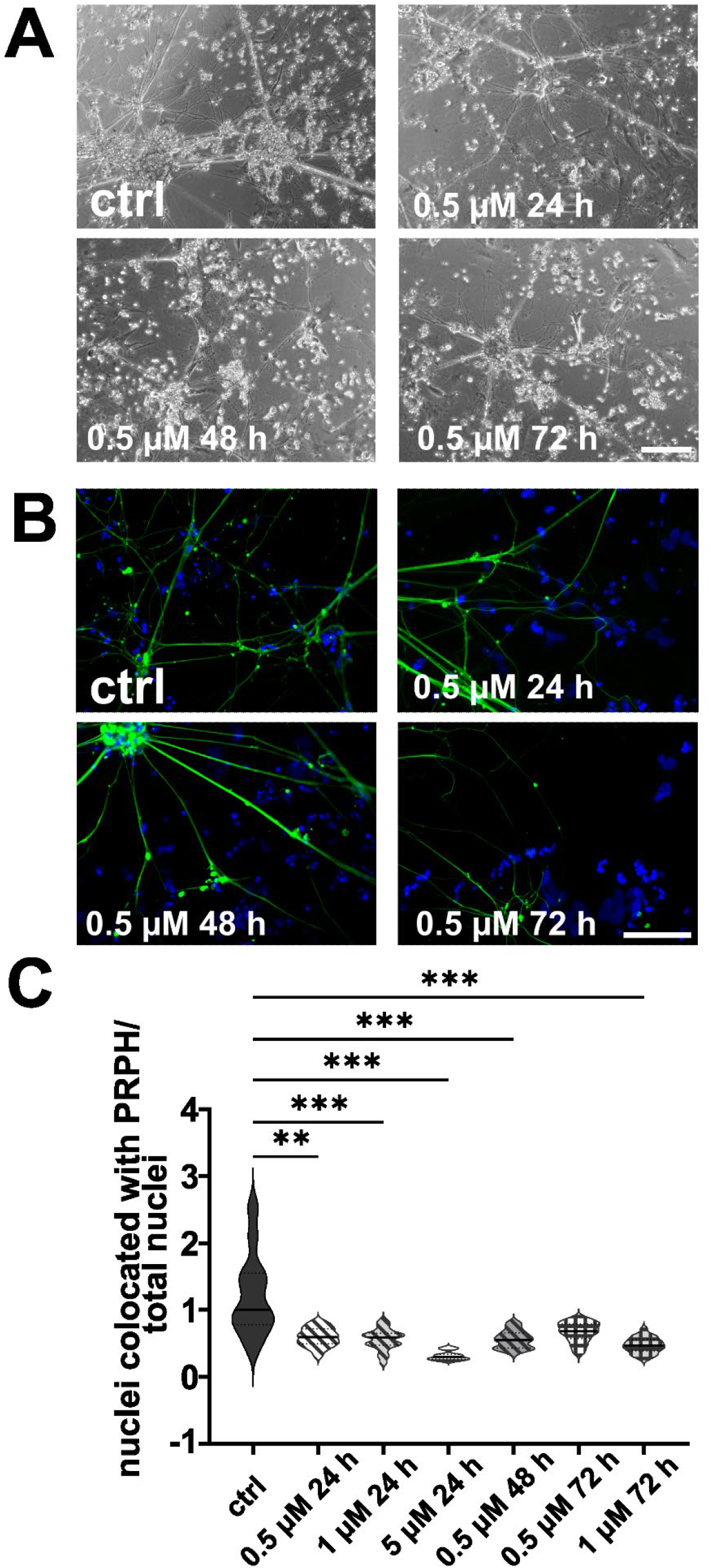
iSN treated with AraC on day +10 of differentiation. Treatment with 0.5 μM AraC as shown in bright field microscopy (A) and ICC photomicrographs did not improve iSN morphology and the number of other cell types regardless of incubation time. ISN-to-total-cell-count ratio was smaller for most conditions where cells survived in comparison to ctrl. 0.5 μM AraC for 72 h was the only condition with similar efficacy as ctrl (C). Data are presented as violin plots, Kruskal-Wallis test with Dunn’s multiple comparison. *p < 0.05; **p < 0.01; ***p < 0.001. Abbreviations: ctrl = control; ICC = immunocytochemistry; PRPH = peripherin. Scale bar = 100 μm.

### Replating iSN shortly after seeding helps reducing non-iSN cells

Cells replated 10, 15, 20, and 30 min after day +10 split showed no difference in morphology (Fig. 5A, B) and iSN-to-total-cell-count ratio (Fig. 5C). When replating was performed 25 min after the initial seeding, the iSN-to-total-cell-count ratio was improved (p < 0.01; Fig. 5C). In general, the replating helped to advance iSN-to-total-cell-count ratio up to 23.88% (Fig. 5D). Replating after 45 min was also tested briefly, but cells had already completely attached at this time point.

**Figure 5:**
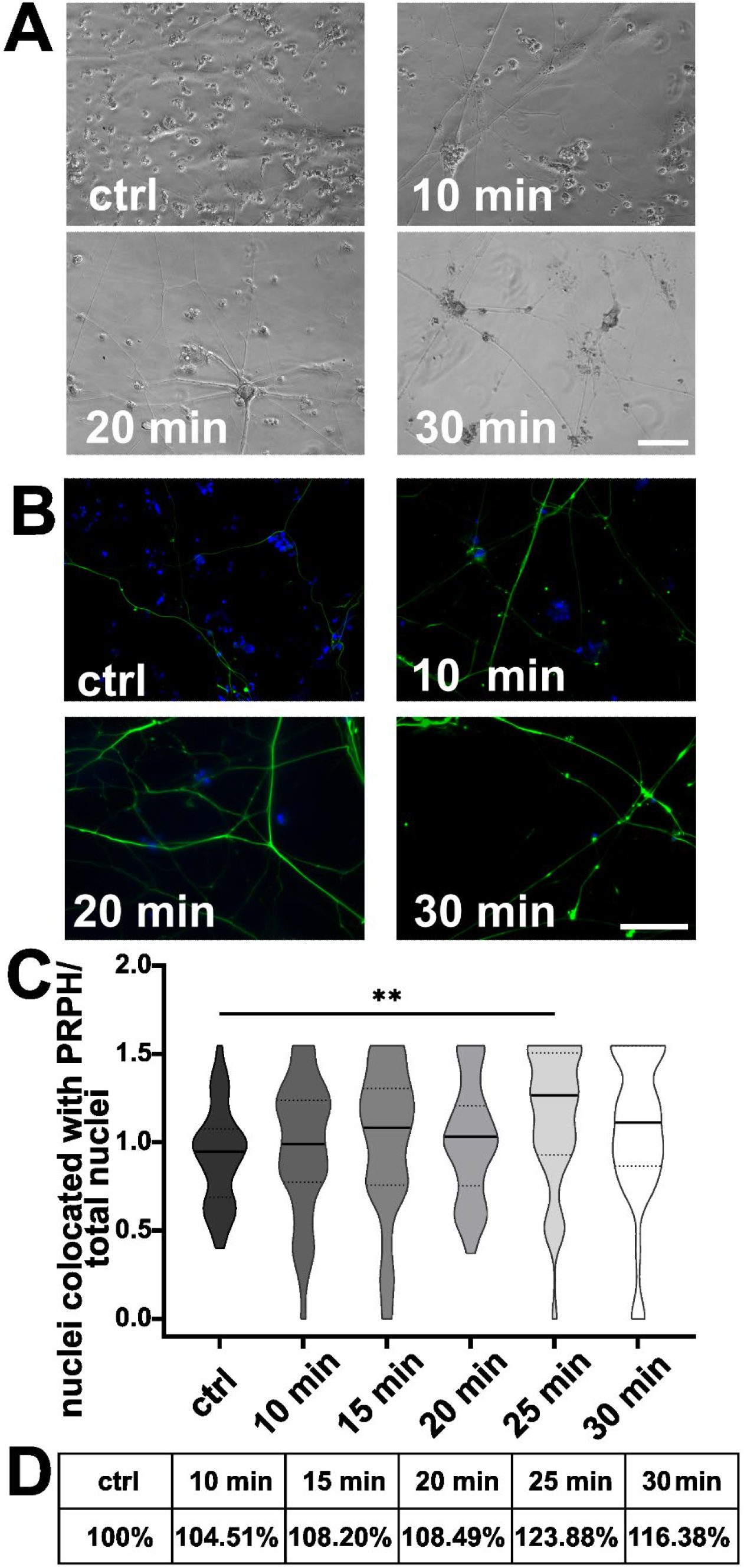
iSN replated onto new CS after day 10 split. Number of non-iSN cells seen in representative bright field microscopy (A) and ICC photomicrographs (B) of cells replated 10, 20 and 30 min after the initial split (additional to 10 μM FdU treatment) was reduced. (C) Replating cells 25 min was most effective, improving iSN-to-total-cell-count ratio compared to the ctrl. (D) Replating generally improved the iSN-to-total-cell-count ratio between 4 and 23.88%. Data are presented as violin plots, Kruskal-Wallis test with Dunn’s multiple comparison. *p < 0.05; **p < 0.01; ***p < 0.001. Abbreviations: ctrl = control; ICC = immunocytochemistry; PRPH = peripherin. Scale bar = 100 μm.

### Long exposure times to FdU and AraC affect cell viability

Cell viability was assessed relative to a control treated with 10 μM FdU for 24 h (Fig. 6A). Treatment of iSN with 0.5 μM AraC for 24 h resulted in an increase in cell viability (p < 0.05). However, extending the exposure to 48 and 72 h at this concentration did not produce relevant changes. Elevating the concentration of AraC to 1 μM and 5 μM had minimal impact on viability, although a marked reduction was noted after three days with 5 μM AraC. While treatment with 10 μM AraC for 24 hours initially increased cell viability, this effect diminished by 48 hours and declined by 72 hours.

**Figure 6:**
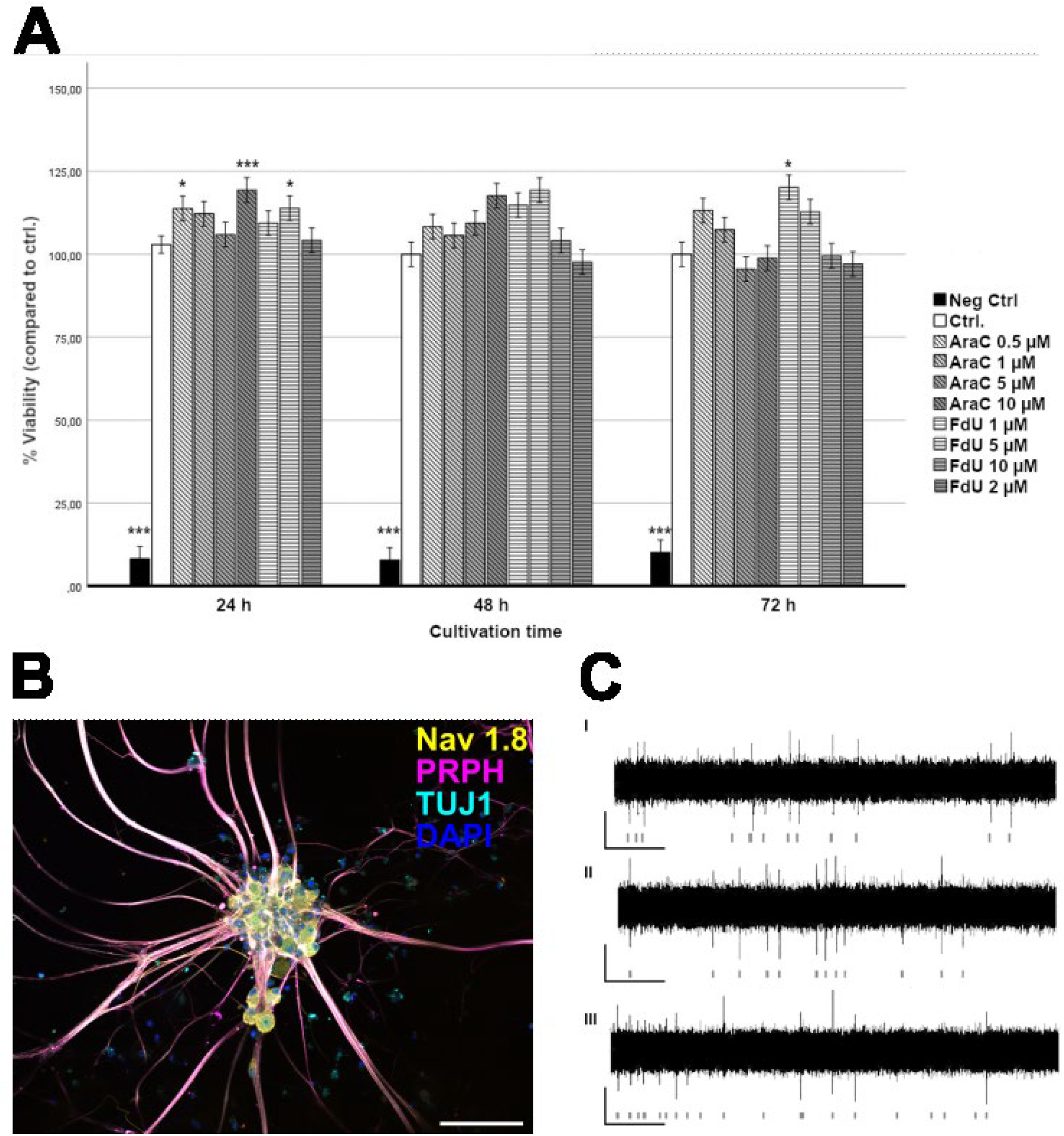
Viability assay, ICC, MEA. (A) Viability assay of iSN following treatment with increasing concentrations of FdU and AraC for up to 72 hours. *p < 0.05; **p < 0.01; ***p < 0.001. (B) iSN generated using our optimized protocol express sensory neuron markers Nav1.8, PRPH and TUJ1. *p < 0.05; **p < 0.01; ***p < 0.001. Abbreviations: ICC = immunocytochemistry; Nav1.8 = voltage-gated sodium channel 1.8, PRPH = peripherin, TUJ1 = Beta-III-tubulin. Scale bar = 100 μm. (C) Spontaneous electrical activity in iSN was measured at 4 (I), 5 (II), and 6 (III) weeks of maturation using a multi-electrode array system, with activity detectable starting from 4 weeks of maturation. Scale: 50 μV/10 sec.

For FdU treatments, concentrations of 1 μM and 5 μM for up to 48 h did not significantly affect viability, except for a slight elevation observed at 72 h with 1 μM FdU and at 24 h with 5 μM FdU. Higher concentrations and longer exposures, particularly 10 μM FdU for 48 and 72 h and 20 μM FdU for 48 and 72 h, resulted in decreased cell viability. No significant changes were detected with 20 μM FdU at 24 h.

### ISN express characteristic markers and are functionally active

Integrity of iSN cultivated under ctrl conditions was confirmed via expression of the sensory neuron markers Nav1.8, PRPH and TUJ1 (Fig. 6B). Electrical activity was observed as early as four weeks into the maturation process and remained present for at least six weeks of maturation, indicating stable development and functionality of iSN (Fig. 6C). Furthermore, the formation of neuronal networks could be observed.

## Discussion

Purity of iSN cultures is often limited by co-existent non-iSN cells which hinder reliable data interpretation and reproducibility. Although several methods have been developed to reduce cellular heterogeneity in neuronal cultures of various origins (Hilgenberg and Smith, 2007; Thirumangalakudi et al., 2009; Irobi et al., 2010; Liu et al., 2013; Schwieger et al., 2016; Clark et al., 2017; Clark et al., 2021; Hirano et al., 2021), no standardized protocol is available (Lampert et al., 2020; Kalia et al., 2024). We have worked on improving cellular purity of iSN using a published protocol (Chambers et al., 2012) and report on the efficacy of cell incubation with 10 μM FdU for 24 h on day +10 split.

The first approach via cell split during differentiation of iSN (Clark et al., 2017) did not increase iSN-to-total-cell count ratios. However, the total number of iSN generated by differentiation including an additional split was substantially higher compared to our ctrl. Hence, early passaging during the differentiation process may ameliorate cell proliferation rates by augmenting the available growth area, but does not influence the number of generated iSN in relation to generated non-iSN.

We could not reproduce the positive effect of MACS twice during the differentiation process increasing culture purity substantially (Hirano et al., 2021). In our hands, MACS did not improve the results compared to 10 μM FdU for 24 h. Also, MACS induces cellular stress and the protocol is time consuming. A single MACS step led to an increased and more homogeneous non-iSN layer in some cases. This might be caused by using neural crest beads for selection, which are not only selective for iSN but also for other neuronal cell types. Thus, we may have generated a more homogeneous culture of other neural crest-derived cells without increasing the number of iSN.

While AraC is often used for animal-derived neuronal cultures (Batista Lobo et al., 2007; Schwieger et al., 2016), it was not feasible in our study. iSN incubated with AraC either showed neuronal blebbing as a sign of stress or did not survive maturation at all. This may be due to the neurotoxic effects of AraC (Baker et al., 1991). AraC also induces apoptosis (Han et al., 2016) and reduces mitochondrial DNA in mouse DRG neurons (Zhuo et al., 2018). Low-dose AraC did not affect iSN, but also allowed growth of non-iSN. In comparison to FdU, AraC was not as efficient. The advantage of FdU over AraC was already shown in hippocampal cell cultures from postnatal rats, where both antimitotic agents were applied to reduce astrocyte contamination (Lesslich et al., 2022).

Furthermore, it was suggested that AraC affects the location of transient receptor potential vanilloid 1 (TRPV1) in human DRG, which should be considered when working with iSN models (Anand et al., 2008). If FdU treatment is not sufficient for downstream experiments and a higher purification is needed, we recommend an additional replating step 25 minutes after the initial split. Replating at later time points was less effective, as most cells had probably already attached.

Treatment with 0.5 μM AraC increased cell viability. High concentrations of FdU (10 μM and 20 μM) for extended treatment times resulted in marked iSN death. Similarly, AraC induced cell death at 5 μM and 10 μM, while lower concentrations (1 μM) were less detrimental, although the overall neuron-to-cell ratio still declined. This suggests that while AraC can be less toxic at lower concentrations, its overall impact on the cellular composition in culture remains important. The observed increase in cell viability in certain treatment groups does likely not reflect iSN health, but rather the proliferation or survival of feeder cells. These cells might be more resilient to the treatments and could overgrow in the culture vessel, skewing viability results.

Eradication of feeder cells was not possible. Furthermore, we observed a decline in neuronal quality and elevated detachment rates in cultures that were almost free of non-iSN. Thus, a certain proportion of non-iSN might be beneficial for neuronal adhesion and providing nutrients to iSN.

ISN generated following our optimized protocol expressed the same sensory neuron markers, PRPH, Nav1.8, and TUJ1, as iSN generated using other protocols (Blanchard et al., 2015; Wainger et al., 2015). Continuing electrical activity, as observed over six weeks, suggests not only the stability of neuronal differentiation but also the maturation of iSN. This prolonged activity likely supports the ongoing synaptic and network refinements critical for functional neuronal network development.

While we assessed many different options to obtain purer iSN cultures, our study has some limitations. First, all experiments were performed using one iPSC line (n = 1 clone). Although inter-individual variation might influence the outcome (Volpato and Webber, 2020), we suggest that the general effects found within this study are transferable to other cell lines. Also, we did not consider how additional modifications in the protocol, e.g., time point of addition and concentration of different small molecules might affect the results (Lampert et al., 2020). The replating was only tested for attachment on glass coverslips coated with bMg, therefore the results and optimal time points may differ when using plastic dishes and other coatings, as these can affect the polarity, arborization, and maturation of cells (Nerli et al., 2020; Setien et al., 2020; Stil et al., 2023).

Still, our work offers a promising strategy for improving iSN purity without compromising cell viability or neuronal quality. However, further research considering inter-individual variability is warranted.

## Data Availability Statement

The original contributions presented in the study are included in the article. For further information, data are available on request from the corresponding author.

## Ethics Statement

The study was approved by the Würzburg Medical Faculty Ethics Committee (#135/15).

## Author Contributions

N.M.S.: Formal analysis, investigation, methodology, visualization, writing – original draft

J.G.: Investigation, methodology, visualization, writing – original draft

F.B.: Investigation, methodology, writing – review and editing

F.K-S.: Investigation, methodology, writing – review and editing

S.O.: Formal analysis, investigation, visualization, writing – review and editing

N.Ü.: Conceptualization, funding acquisition, project administration, supervision, writing – original draft

## Acknowledgments

We thank our undergraduate student Simran Jeet Kaur (Department of Neurology, University Hospital Würzburg, Germany) for help during cell cultivation.

